# In-field genetic stock identification of overwintering coho salmon in the Gulf of Alaska: Evaluation of Nanopore sequencing for remote real-time deployment

**DOI:** 10.1101/2021.05.27.445905

**Authors:** Christoph M. Deeg, Ben J. G. Sutherland, Tobi J. Ming, Colin Wallace, Kim Jonsen, Kelsey L. Flynn, Eric B. Rondeau, Terry D. Beacham, Kristina M. Miller

## Abstract

Genetic stock identification (GSI) by single nucleotide polymorphism (SNP) sequencing has become the gold standard for stock identification in Pacific salmon, which are found in mixed-stocks during the oceanic phase of their lifecycle. Sequencing platforms currently applied require large batch sizes and multi-day processing in specialized facilities to perform genotyping by the thousands. However, recent advances in third-generation single-molecule sequencing platforms, like the Oxford Nanopore minION, provide base calling on portable, pocket-sized sequencers and hold promise for the application of real-time, in-field stock identification on variable batch sizes. Here we report and evaluate utility and comparability of at-sea stock identification of coho salmon *Oncorhynchus kisutch* based on targeted SNP amplicon sequencing on the minION platform during the International Year of the Salmon Signature Expedition to the Gulf of Alaska in the winter of 2019. Long read sequencers are not optimized for short amplicons, therefore we concatenate amplicons to increase coverage and throughput. Nanopore sequencing at-sea yielded stock assignment for 50 of the 80 assessed individuals. Nanopore-based SNP calls agreed with Ion Torrent based genotypes in 83.25%, but assignment of individuals to stock of origin only agreed in 61.5% of individuals highlighting inherent challenges of Nanopore sequencing, such as resolution of homopolymer tracts and indels. However, poor representation of assayed coho salmon in the queried baseline dataset contributed to poor assignment confidence on both platforms. Future improvements will focus on lowering turnaround time, accuracy, throughput, and cost, as well as augmentation of the existing baselines, specifically in stocks from coastal northern BC and Alaska. If successfully implemented, Nanopore sequencing will provide an alternative method to the large-scale laboratory approach. Genotyping by amplicon sequencing in the hands of diverse stakeholders could inform management decisions over a broad expanse of the coast by allowing the analysis of small batches in remote areas in near real-time.

## Introduction

The semelparous and anadromous life history of Pacific salmon makes them crucial to coastal and terrestrial ecosystems around the North Pacific by connecting oceanic and terrestrial food webs and nutrient cycles (Cederholm et al. 1999). Salmon are highly valued by the northern Pacific Rim nations due to their significant contribution to commercial and recreational fishing harvests as well as their cultural importance, especially amongst Indigenous peoples (Lichatowich 2001). Despite this significance, many wild Pacific salmon stocks have experienced population declines due to a combination of compounding factors such as overexploitation, spawning habitat alterations, pathogens and predators, prey availability, and climate change (Miller et al. 2014). Efforts to rebuild stocks include habitat restoration, artificial stock enhancements, as well as stock specific monitoring through a number of assessment methods to inform targeted management strategies (Hinch et al. 2012). These monitoring strategies include spawning escapement and smolt survival assessments as well as test fisheries in riverine and coastal waters (Woodey 1987). While these tools allow the assessment of health and productivity of individual stocks and therefore the targeted management of individual populations in North American coastal and riverine environments, the open ocean phase of the life cycle of Pacific salmon remains poorly studied despite the observed large temporal shifts in marine survival over recent decades (Holtby, Andersen, and Kadowaki 1990). Compounding this issue, Pacific salmon are usually found in mixed-stock schools in the ocean, meaning that fish from home streams as distant as North America and Asia might be found in the same school, making stock-specific management challenging (Wood, Rutherford, and McKinnell 1989).

To overcome the challenges of mixed-stock management, stock identification has in the distant past utilized characteristic scale and parasite patterns, as well as the marking of hatchery-enhanced fish by coded-wire tagging (Wood, Rutherford, and McKinnell 1989; Cook and Guthrie 1987; Jefferts, Bergman, and Fiscus 1963). More recently, genetic stock identification (GSI) using minisatellite, microsatellite, and ultimately single nucleotide polymorphisms (SNPs) as markers has proven superior in delivering high-throughput insights into the stock composition of salmon (Beacham et al. 2017, 2018; Miller, Withler, and Beacham 1996). Specifically, the large baseline of population-specific SNP frequencies and targeted amplification of such loci now allow for unprecedented resolution of stock origin in many species of salmon at reduced biases (Beacham et al. 2018, 2017; Ozerov et al. 2013; Gilbey et al. 2017). However, current sequencing approaches, based on second generation sequencing platforms (e.g. illumina and Ion Torrent), mean that only sequencing large batches of individuals, known as “genotyping by the thousands” (GT-seq), is economically and practically feasible (Beacham et al. 2017, 2018; Campbell, Harmon, and Narum 2015). These approaches require a specialized laboratory and several days turnover for the library preparation and sequencing, even under highly automated settings. These constraints limit the utility of SNP-based GSI for spatially or temporally restricted assessments because samples need to be transported to the laboratory for analysis, as has been the case for all GSI methods to date. Specifically, for time-sensitive stock-specific harvest management decisions, an in-field real-time SNP-based GSI approach with greater flexibility in sample batch size would be desirable.

Recent advances in third-generation single-molecule sequencing platforms like the Oxford Nanopore minION allow real-time sequencing on a pocket-sized portable sequencer that requires little library preparation, therefore enabling sequencing in remote locations (Mikheyev and Tin 2014; Quick et al. 2016). However, several technical hurdles to adapting Nanopore sequencing to SNP GSI exist. While Nanopore sequencing can yield extremely long reads, the number of sequencing pores and their loading is limited, resulting in low throughput when sequencing short reads, such as amplicons. An additional problem is the relatively high error rate inherent to this novel technology. Since the SNP GSI protocols are based on the amplification of short amplicons via targeted multiplex PCR, sequencing throughput of such short amplicons on the Nanopore platform is comparatively low, as the number of sequencing pores is the rate limiting factor. This is especially problematic since high coverage is needed to compensate for the higher error rate of Nanopore generated sequences. A promising approach to overcome these limitations is the concatenation of PCR amplicons that allows the sequencing of several amplicons within a single read, thereby exponentially increasing throughput for genotyping (Cornelis et al. 2017; Schlecht et al. 2017).

Here, we report on the development and performance of a novel Nanopore-based in-field SNP GSI method by adapting existing SNP GSI technology to the Nanopore platform using a concatenation approach (Schlecht et al. 2017). We aim to demonstrate in-field feasibility, repeatability and comparability to established platforms. As a proof of concept, in-field stock ID was performed in the Gulf of Alaska onboard the research vessel *Professor Kaganovsky* during the IYS expedition in February and March of 2019.

## Materials and Methods

### Field Lab equipment and workspace

The field equipment onboard the *Professor Kaganovsky* research trawler consisted of a PCR thermocycler, a mini-plate centrifuge, a microcentrifuge, a Qubit fluorimeter (Thermo Fisher), a vortexer, a minION sequencer, a laptop with an Ubuntu operating system (Ubuntu v.14.06), as well as assorted pipettes and associated consumables like filter tips (Figure 1). The required infrastructure onboard included a 4°C fridge, a −20°C freezer, a power supply, as well as a physical workspace. The entire equipment configuration required was under $10,000 CAD.

**Figure 1:**
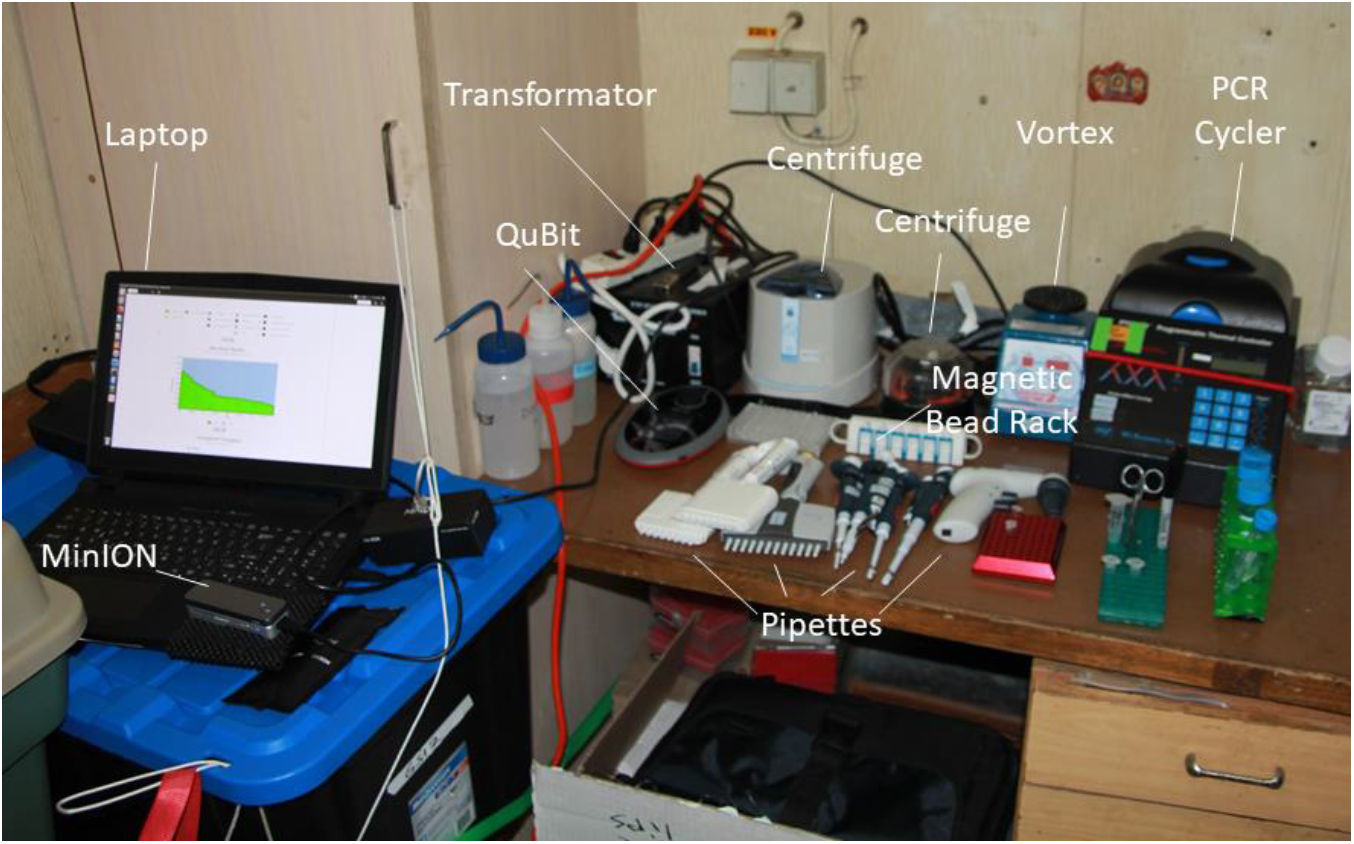
*Workspace abroad the* Professor Kaganovsky *vessel during the International Year of the salmon signature expedition*.

### Tissue sample collection and DNA extraction

Salmon were captured by the research trawler *Professor Kaganovsky* during the 2019 International Year of the Salmon (IYS) Signature expedition in the Gulf of Alaska (Figure 2). We collected fin clips of coho salmon (*Oncorhynchus kisutch*) and froze them individually until DNA extraction, or immediately processed once a suitable batch size had been accumulated. DNA extraction from 2 *x* 2 *x* 2mm fin-tissue clips was performed in a 96-well PCR plate using 100μl of QuickExtract solution (Lucigen, USA) according to the manufacturer’s instructions.

**Figure 2:**
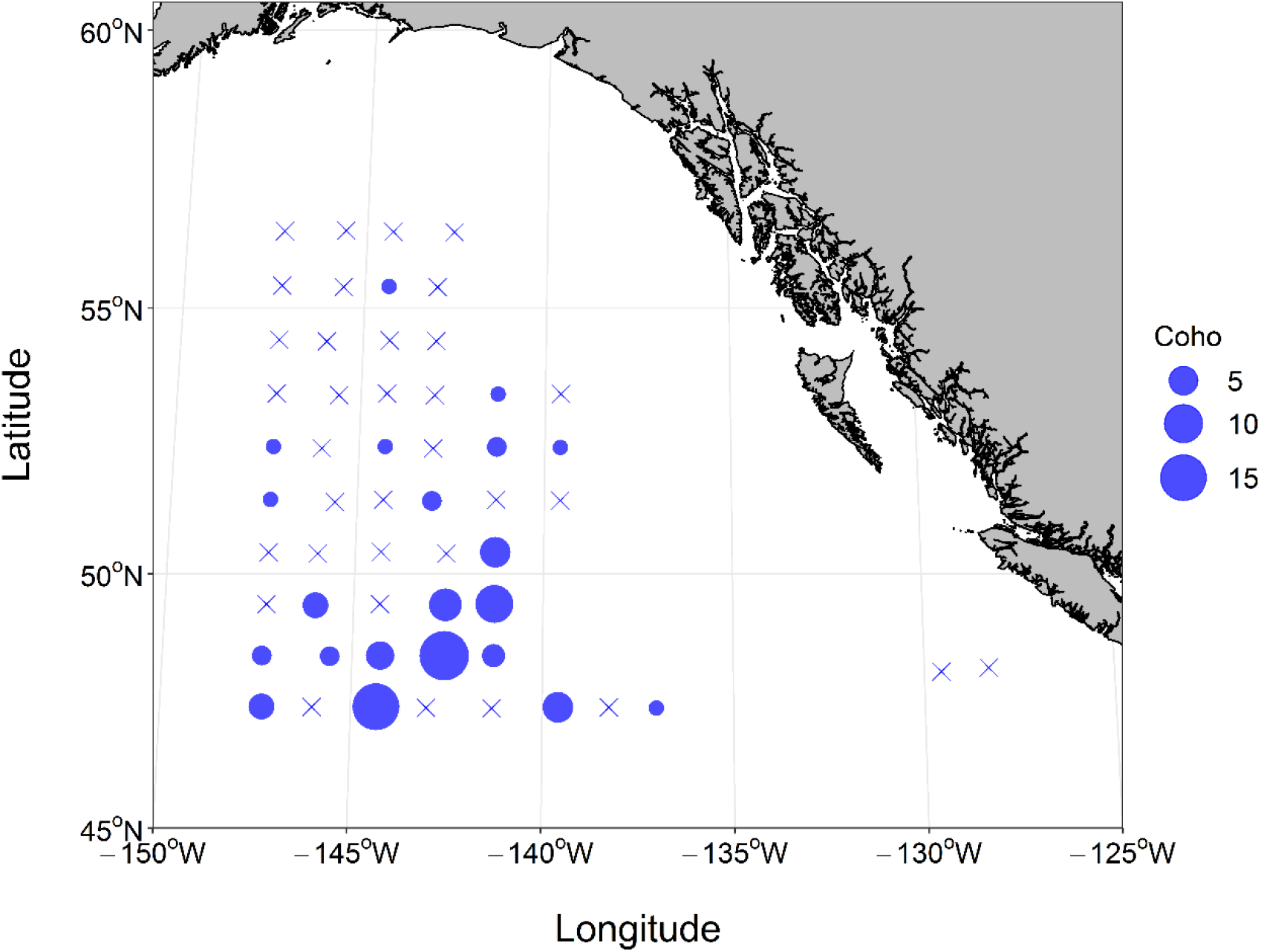
Coho salmon capture numbers and location during the International Year of the Salmon Gulf of Alaska expedition.Size of circles corresponds to catch size, X indicates a trawl without coho capture.

### Multiplex PCR and Barcoding

Multiplex PCR with a custom panel of primers targeting 299 loci of known SNPs was performed using 0.25μl of DNA extract as template using the AgriSeq HTS Library Kit Amplification Mix PCR mastermix (ThermoFisher) in a 10μl reaction according to Beacham et al. (Beacham et al. 2017). Next,we prepared amplicons for ligation by end-prepping amplified strands with AgriSeq HTS Library Kit Pre-ligation Enzyme. ONT barcode adapters (PCR Barcoding Expansion 1-96, EXP-PBC096, Oxford Nanopore Technologies, UK) were then ligated to the amplicons by blunt-end ligation with the Barcoding Enzyme/Buffer of the AgriSeq HTS Library Kit according to manufacturer’s instruction. After bead-cleanup (1.2:1 bead:sample, AMPure XP beads, Beckman Coulter, USA) we added the ligation products, barcodes and barcoding adapters (PCR Barcoding Expansion 1-96, EXP-PBC096, Oxford Nanopore Technologies, UK) by PCR using Q5 polymerase mastermix (NEB, USA) for individual fish identification according to manufacturer’s protocol in a 25μl reaction (98°C for 3 min; 25 cycles of 98°C for 10 s, 70°C for 10 s, 72°C for 25 s; 72°C for 2 min). Barcoded libraries were then pooled and cleaned using 1.2:1 bead cleanup, before DNA yield of a subset of samples (12.5%) was analyzed by Qubit (dsDNA HS Assay Kit, ThermoFisher, USA).

### Amplicon concatenation

To improve throughput on the minION, we concatenated amplicons using inverse complementary adapters (Figure 3). After end prep using Ultra II End Repair/dA-Tailing Module Module (NEB, USA), the library was split into two equal volume subsets. Custom inverse complementary adapters that had inverse complementary terminal modifications to ensure unidirectional ligation (3’-T overhang and 5’ phosphorylation) were ligated onto both ends of the respective subsets using the Ultra II Ligation Module (NEB, USA) according to manufacturer’s instructions and purified with 1:1 bead cleanup (Figure 3). The custom adapters were adapted from Schlecht et al (Schlecht et al. 2017): Adapter A: 5’P-ACAGCGAGTTATCTACAGGTTCTTCAATGT + ACATTGAAGAACCTGTAGATAACTCGCTGTT; Adapter B: 5’P-ACATTGAAGAACCTGTAGATAACTCGCTGT + ACAGCGAGTTATCTACAGGTTCTTCAATGTT). Amplicons with adapters added to them were subsequently amplified again with a single primer (5μl; ACATTGAAGAACCTGTAGATAACTCGCTGTT for adapter A, ACAGCGAGTTATCTACAGGTTCTTCAATGTT for adapter B) in 25μl Q5 reactions according to manufacturer’s instructions with the following thermal regime: 98°C for 3 min; 30 cycles of 98°C for 10 s, 68°C for 15 s, 72°C for 20 s; 72°C for 2 min). After 1:1 bead cleanup, we pooled both subsets in equimolar ratios after Qubit quantification to verify both reactions worked, and then subjected to a primer-free, PCR-like concatenation due to heterodimer annealing and elongation in 25μl Q5 reaction, using the complementary adapter sequence ligated onto the amplicons as primers cycled under the following thermal regime: 3 cycles of 98°C for 10 s, 68°C for 30 s, 72°C for 20 s; followed by 3 cycles of 98°C for 10 s, 68°C for 30 s, 72°C for 30 s; followed by 3 cycles of 98°C for 10 s, 68°C for 30s,72°C for 40s; followed by 3 cycles of 98°C for 10 s, 68°C for 30 s, 72°C for 50s; and finally followed by 72°C for 2 min (Figure 3).

**Figure 3:**
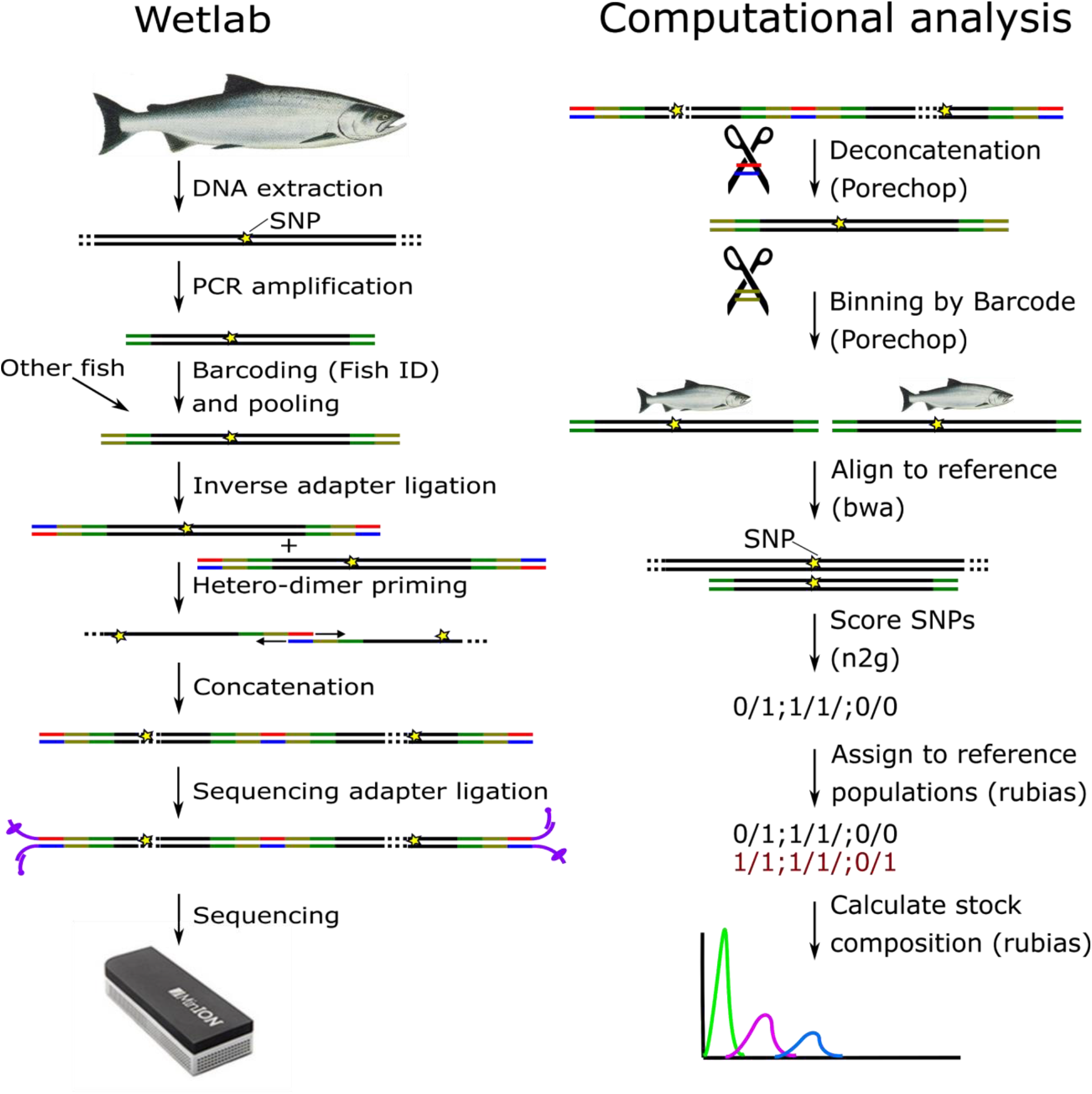
Simplified wet-lab workflow for DNA extraction, amplification, barcoding, and concatenation before sequencing and pipeline of the following computational analysis. DNA is shown in black, amplification primers in green, fish ID barcodes in olive, concatenation adapters in red/blue, and sequencing adapters in purple.

### Library Preparation and sequencing

The concatenated amplicons were prepared for Nanopore sequencing using the ONT Ligation Sequencing Kit (LSK109) according to the manufacturer’s instruction. In brief, after end-prep using the Ultra II Endprep Module and bead cleanup, we ligated proprietary ONT sequencing adapters onto the concatenation adapters by blunt-end ligation using the proprietary ONT Buffer and the TA quick ligase (NEB, USA; note: this standard sequencing step not shown in Figure 3). After additional bead-cleanup and washing with the short fragment buffer (SFB: ONT, UK) according to the manufacturer’s protocol,we loaded the library onto a freshly primed flow cell (MIN 106 R9.4.1: ONT, UK) according to the manufacturer’s instruction.

### Nanopore sequencing

After flow cell priming and loading of the library, the flow cell was placed on the minION sequencer. Sequencing and basecalling into fast5 and fastq was performed simultaneously using minKNOW (version 3.1.8) on an Ubuntu 14.06 platform.

### Deconcatenation and binning

First, all fastq raw reads that passed default quality control in minKNOW were combined into bins of 500k reads each. This had empirically been determined to be the maximum number of reads allowing simultaneous processing in the downstream analysis on our platform (Ubuntu 14.06, 31.2 GiB RAM 7700K CPU @ 4.20GHz × 8). Reads containing concatenated amplicons were deconcatenated and the concatenation adapter sequence was trimmed off the remaining sequence using porechop (https://github.com/rrwick/Porechop) with a custom adapter file (“adapters.py”) that only contained the concatenation adapter under the following settings: porechop-runner.py -i input_raw_reads.fastq -o output/dir -t 16 --middle_threshold 75 --min_split_read_size 100 --extra_middle_trim_bad_side 0 --extra_middle_trim_good_side 0

We binned the deconcatenated reads by barcode corresponding to fish individuals by using porechop with the provided default adapters file and the following settings: porechop-runner.py -i input_deconcatenated_reads.fastq -b binning/dir -t 16 --adapter_threshold 90 -- end_threshold 75 --check_reads 100000

After this step, all reads from the corresponding barcode bins corresponding to the same individual across the different 500k sub-bins were combined for downstream analysis. See https://github.com/bensutherland/nano2geno/ for source scripts for analysis.

### Alignment and SNP calling

We aligned the binned reads to the reference amplicon sequences using BWA-MEM and indexed using samtools (Li et al. 2009; Li and Durbin 2009). Alignment statistics for all loci were generate using pysamstats (https://github.com/alimanfoo/pysamstats; flags: -t variation -f) and we extracted the nucleotides observed at the relevant SNP hotspot loci from the resulting file using a custom R script by looping through the results file guided by a SNP location file. Finally, we compared the observed nucleotide distributions at SNP hotspots with to the hotspot reference and variant nucleotides and scored as homozygous reference when ≥66% of the nucleotides were the reference allele, heterozygous when the reference allele was present <66% and the variant allele > 33%, or as homozygous variant (when the nucleotides were ≥66% the variant allele) using a custom R script to generate a numerical locus table. We visually inspected alignments determined to be problematic using the IGV viewer (Robinson et al. 2011). The full pipeline titled “nano2geno” (n2g) including all custom scripts can be found at https://github.com/bensutherland/nano2geno/ (Figure 3).

### Mixed-stock Analysis

We performed mixture compositions and individual assignments using the R package rubias (Moran and Anderson 2019) with default parameters against the coho coastwide baseline of known allele frequencies for these markers established by Beacham et al.

### Ion torrent sequencing

To confirm the results obtained by Nanopore sequencing, the tissue samples were sequenced using an Ion Torrent sequencer according to Beacham et al. 2017 (Beacham et al. 2017). In brief: DNA was extracted from the frozen tissue samples using Biosprint 96 SRC Tissue extraction kit, and multiplex PCR and barcoding with Ion Torrent Ion Codes was performed using the AgriSeq HTS Library Kit (ThermoFisher). The libraries were then prepared with the Ion Chef for sequencing on the Ion Torrent Proton Sequencer and SNP variants were either called by the Proton VariantCaller (ThermoFisher; Torrent Suite 5.14.0) software or the custom SNP calling script of the nano2geno pipeline. The resulting locus score table was then analyzed using rubias as described above.

### Concordance assessment

We assessed concordance between sequencing platforms on SNP level. A PCoA analysis was performed using the R package ape based on a reference vs allele call matrix (Paradis and Schliep 2019). Additionally, calls (reference vs. alternate allele) were compared for each sample and marker individually, then averaged by individual, and then averaged by the entire assessed population. Similarly, we compared stock assignment by rubias by comparing the reporting unit or collection as assigned and scoring a match (1) or non-match (0). These scores were then averaged again to generate the final concordance or repeatability score as a percentage.

## Results

### In-field Nanopore Sequencing

During the International Year of the Salmon Signature expedition to the Gulf of Alaska in February and March 2019, in-field single nucleotide polymorphism genetic stock identification (SNP GSI) was performed on coho salmon as the tissues became available. A total of 75 coho salmon were analyzed in two sequencing runs at different points during the expedition, representing 77% of all coho salmon captured during the expedition. The first sequencing run was performed on February 26^th^ and included 31 individuals. Library preparation onboard the vessel took 14h. However, faulty flow cell priming resulted in only approximately half the detected pores being active (843 pores) in this first attempt. Of these pores, no more than 25% were actively sequencing at any time, highlighting the challenges of utilizing sensitive equipment under field conditions including excessive ship movement. Accordingly, sequencing for 30h and base-calling for 34h resulted in only 1.44M reads, 49% of which passed quality control. The read length distribution showed several large concatenated amplicons up to 7,095 bp with a mean length of 825 bp (Figure 4). Deconcatenation resulted in a read inflation by a factor of 2x (702k to 1,444k reads). After binning, reads per individual ranged from 1,983 to 86,467 reads with a mean of 13,709 reads (SD: 15,370), and 722,174 reads that were not able to be assigned (50% of total deconcatenated reads) (Figure 4, Figure 5, Figure 6).

**Figure 4:**
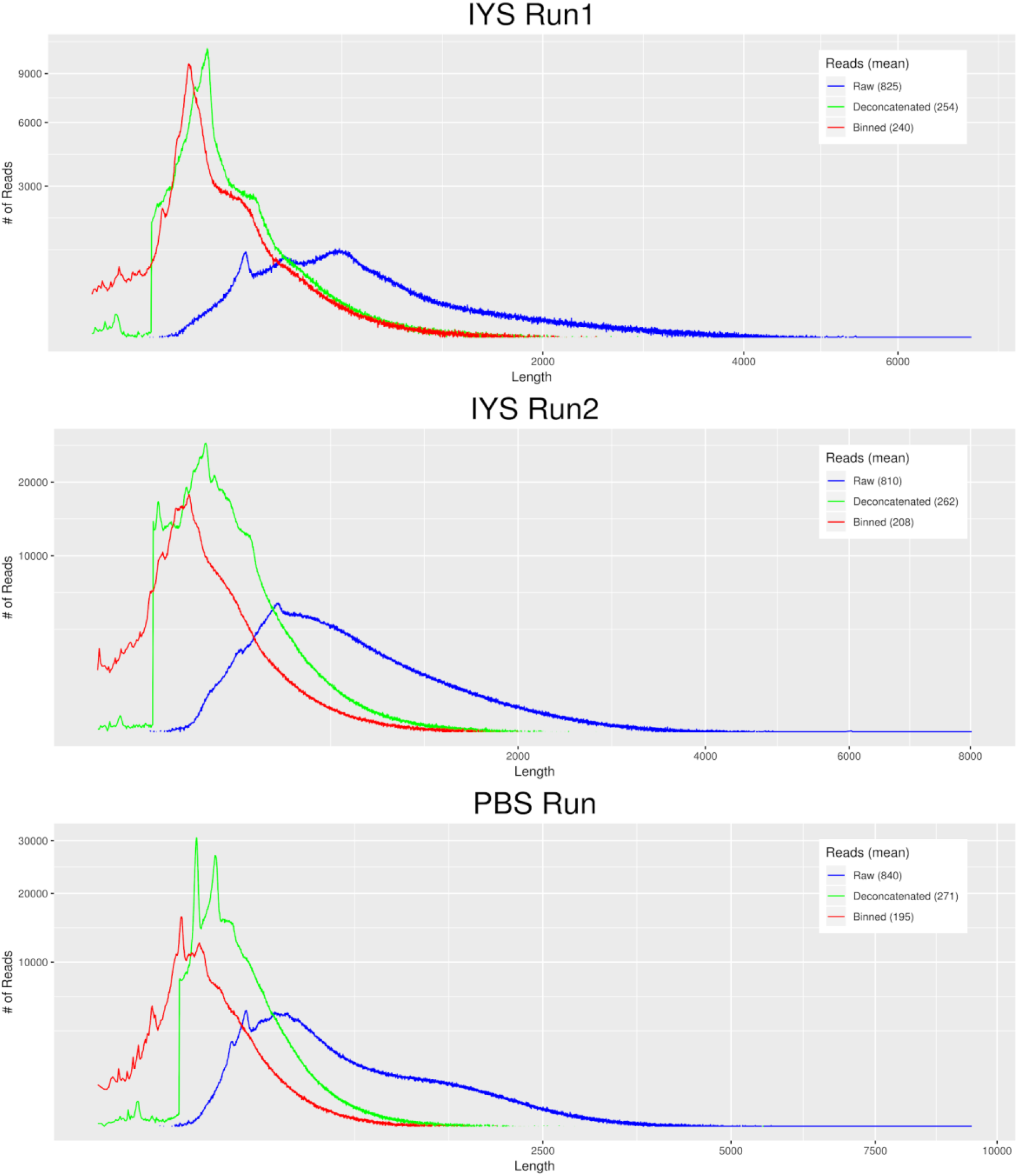
Read length distribution of sequences by Nanopore run: Raw concatenated Nanopore reads shown in red, deconcatenated reads in blue and binned reads in green with the mean read length shown in the legend in parentheses. IYS Run 1 and IYS Run 2 were performed at-sea onboard the *Professor Kaganovsky*, the control run upon return from the expedition is titled “PBS Run”. All axes are square-root transformed.

**Figure 5:**
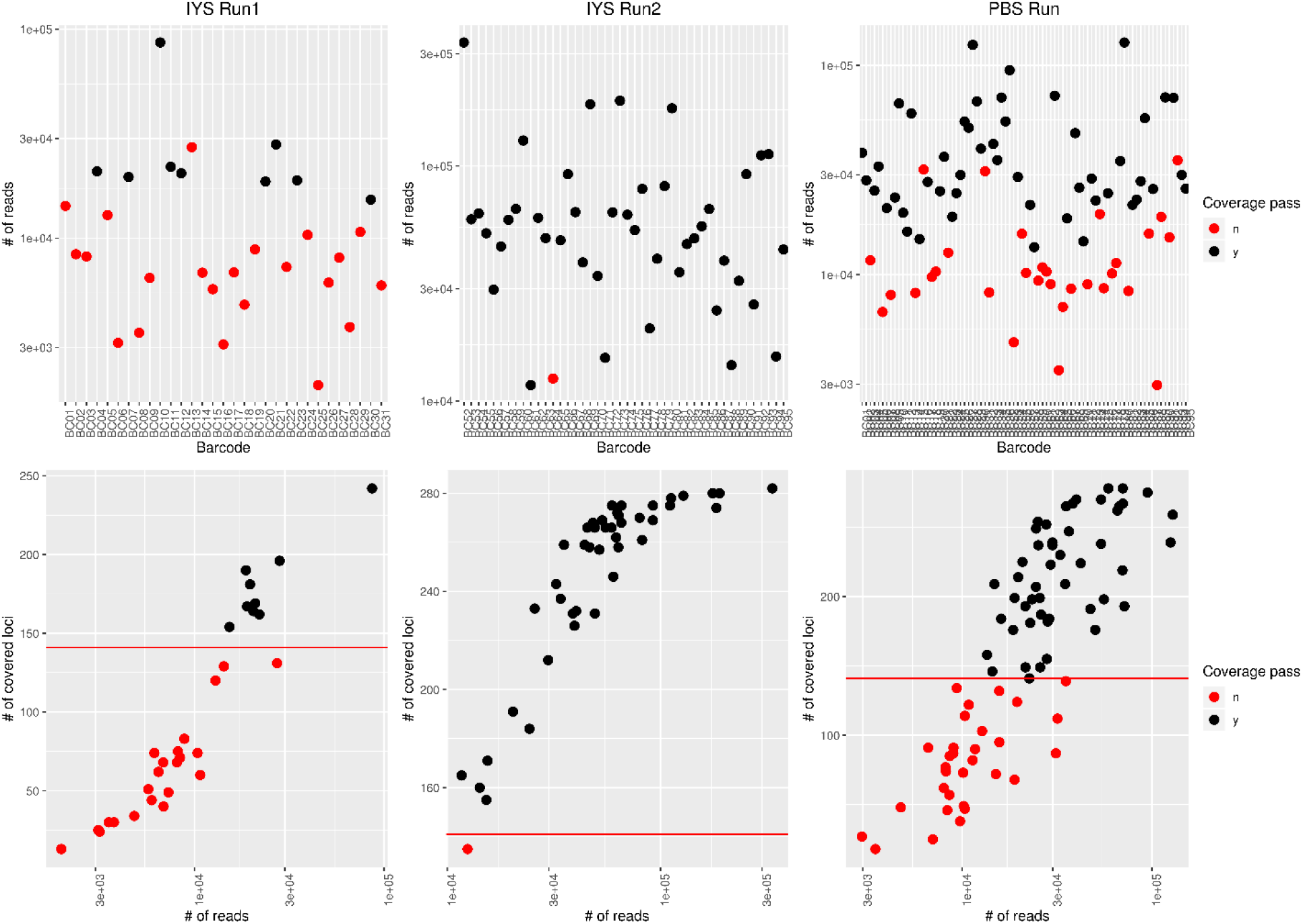
Nanopore sequencing statistics. The top row shows the number of reads binned by individual (barcode). The bottom row shows the number of loci with sufficient read depth (≥10 reads) by the number of reads for each sample. The threshold for downstream analysis was set to 141 loci (50% of loci for this panel) and is indicated in the red line in the second row. Individuals (barcode bins) are coloured according to whether they passed this coverage threshold (black) or not (red).

**Figure 6:**
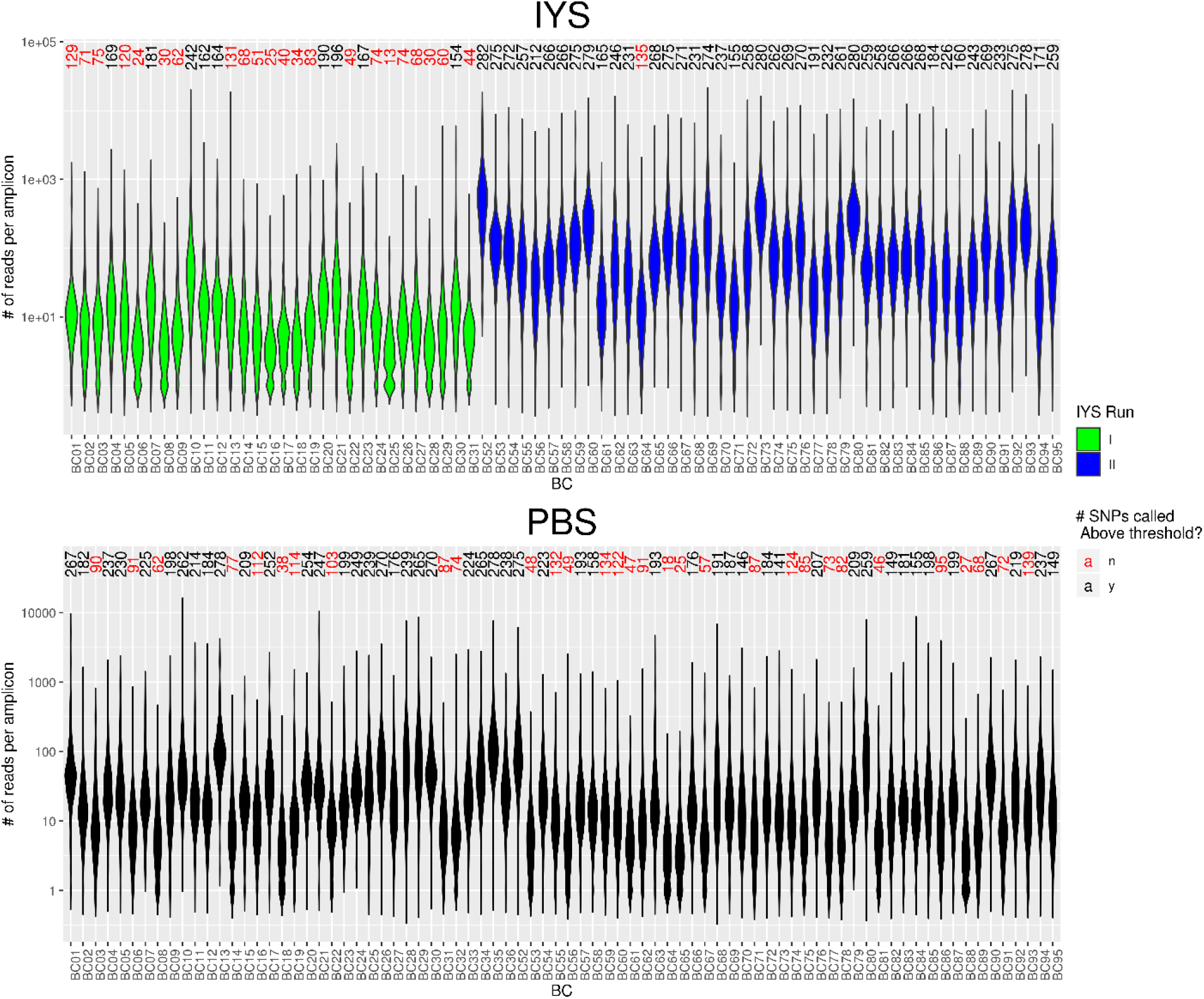
Number of reads per amplicon per individual (barcode) of Nanopore sequencing runs. The violin plot shows the distribution of number of reads assigned to unique SNP-containing amplicons within an individual. Green and blue colors denote the two separate sequencing runs during the IYS expedition (top), and black indicates the run at the laboratory (PBS; bottom). *Above each individual violin plot is the total number of amplicons for that individual for which sufficient reads were present to call the genotype, color indicates if enough amplicons were called for downstream analysis (black) or not* (*red*). The order of individuals is matched in the top and bottom plots.

The second sequencing run was performed on March 10^th^ 2019 with 44 coho salmon. Library preparation again took 14h and sequencing on a new flow cell took 15h, starting with 1,502 available pores, and up to 65% actively sequencing pores, and resulted in 4.48M reads, 76% of which passed quality control. Read lengths averaged 810 bp with a maximum length of 8,023 bp (Figure 4). Due to the large number of reads and the limited power of the computer being used for the analysis, base-calling into fastq took three days. Deconcatenation resulted in a read inflation of a factor of 1.7x (3.4M to 5.8M) (Figure 4). Reads per individual showed a mean of 67,636 reads (SD: 59,393; min: 11,684; max: 335,348), with 722,179 reads remaining unassigned (12%) (Figure 4, Figure 5, Figure 6).

Upon return from the expedition, we sequenced 80 individuals, including all those previously genotyped aboard the vessel, in a single MinION run using the expedition setup starting from the frozen tissues from the expedition. We sequenced for 42h to maximize the total number of reads with 60% of 2,048 available pores actively sequencing resulting in 5.32 M reads. Of these reads, 3.20 M passed quality control. Again, large concatenated amplicons up to 9,449 kb were observed, with a mean read length of 840 bp, and deconcatenation resulted in 4.54 M reads (1.4x inflation) (Figure 4). The mean number of reads per bin was 29,439 (SD: 25,000) and ranged from 2,969 to 128,718 reads per individual, with 1,413,626 unassigned reads (31%) (Figures 4–6).

Despite the absence of normalization between samples prior to multiplex PCR, barcoding, and loading, the binning distribution across samples was relatively even with only a few apparent outliers observed (Figure 5, Figure 6). The minimum number of reads per individual sample necessary to cover sufficient loci (at a minimum depth of 10 sequences per locus) for downstream stock assignments (i.e., at least 141 loci per sample) is around 2,000 reads (Figure 5, Figure 6).

### Nanopore sequencing data requires loci reassessment for efficient SNP calling

After alignment to the reference sequences for SNP calling, Nanopore sequence data showed a comparatively higher error rate than Ion Torrent reads, as expected, with abundant indels that frequently led to lower alignment scores than those obtained by the Ion Torrent data (Ion Torrent average alignment score: 25.6 MAPQ; Nanopore average alignment score: 13.9 MAPQ). Specifically, regions containing homopolymer tracts were poorly resolved, as had previously been reported (Cornelis et al. 2017). Several instances could be identified where the homopolymer presence near the SNP locus caused problematic alignments and therefore resulted in SNP calls not matching those found by the Ion Torrent on the same individual (Figure 7). Accordingly, six such loci were excluded from downstream analysis (Supp. Table 1). Other loci were excluded from the analysis due to absence of coverage (four loci) or the inability of the custom n2g pipeline to call MNPs (multi-nucleotide polymorphisms) or deletions (seven loci), bringing the number of accessed loci from 299 to 282 loci. Other loci showing apparent differences between Nanopore and Ion Torrent sequence data (n = 21) were retained as no apparent explanation for the discrepancies could be identified.

**Figure 7:**
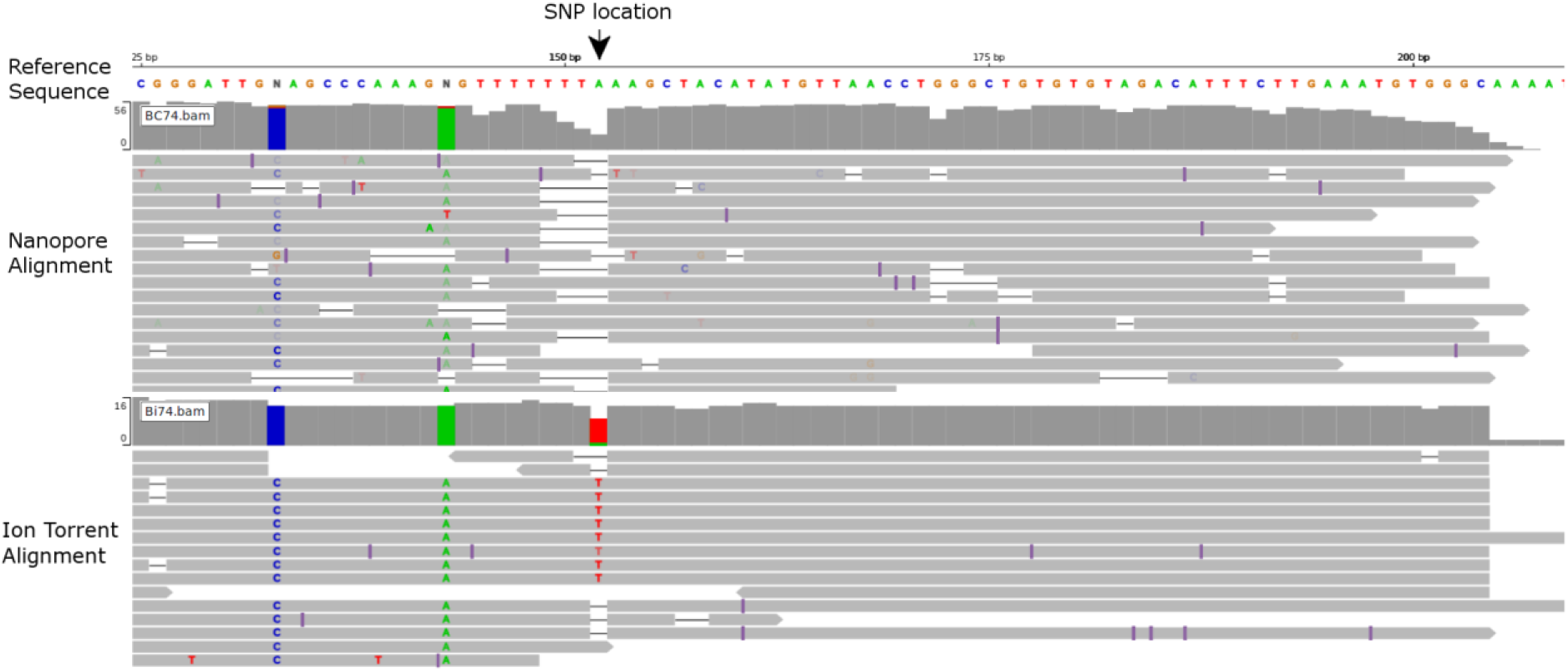
Comparison of sequence alignment of Nanopore and Ion Torrent sequences from the same individual against a SNP locus preceded by a homopolymer tract. Nanopore sequences show a higher number of indels, specifically associated with the poly-T homopolymer tract (145-151bp) directly preceding the SNP location (152bp). Alignment was visualized here using IGV (Robinson et al. 2011)

After the removal of the discrepancies due to MNP, homopolymer, or deletion presence, the SNP cutoff for downstream analysis was set to 141 loci (50%). Only nine of 31 individuals (29%) of the first IYS sequencing run with problematic flow cell priming passed this threshold. In the second IYS sequencing run, 43 of 44 individuals passed the threshold (98%). The repeat run performed at the Pacific Biological Station resulted in 50 of the 80 (63%) that passed this threshold (Figure 6).

### Platform biases lead to moderately altered SNP calling compared to Ion Torrent sequencing

To assess the discrepancies between sequencing platforms, individuals that passed the genotyping rate threshold of 141 called loci (50% genotyping rate) in all data sets (i.e., Nanopore data during the expedition analyzed with n2g: “nano IYS”, Nanopore acquired during the repeat run upon return from the expedition, analyzed with n2g: “nano PBS”, Ion Torrent sequencing data analyzed with variant caller: “ion vc”, Ion Torrent analyzed with n2g: “ion n2g”) were included in a PCoA analysis on the SNP genotypes (Figure 8). This comparison excluded the MNP, deletion, and homopolymer loci (see above), but retained those without an explanation as to why the genotyping did not match. However, there was still an apparent separation by sequencing platform across the highest-scoring dimension (Figure 8). This trend was reflected by 83.9% of SNP calls generated by Nanopore sequencing during the IYS expedition (nano IYS) and 83.7% of SNP calls generated during the repeat run upon return (nano PBS) matching the SNP calls based on Ion Torrent data (ion n2g). The agreement on SNP call between both Nanopore runs (comparing reference or alternate scores for both alleles from nano IYS vs nano PBS) was 84.4%, highlighting the inter run variability associated with current Nanopore sequencing. There was a slight correlation observed between the number of Nanopore reads per individual and the concordance with Ion Torrent SNP calls, suggesting that read depth is only a minor factor influencing SNP call concordance at the current threshold of a minimal alignment depth of 10x per site for Nanopore reads (Figure 9). Excluding MNPs, deletions, and homopolymer issues, the influence of the SNP calling pipeline (n2g vs. variant caller) appears negligible compared to the differences by sequencing platform (Figure 8). Accordingly, SNPs scored based on the same Ion Torrent data sequence matched in 99.21% of cases between the two genotyping pipelines.

**Figure 8:**
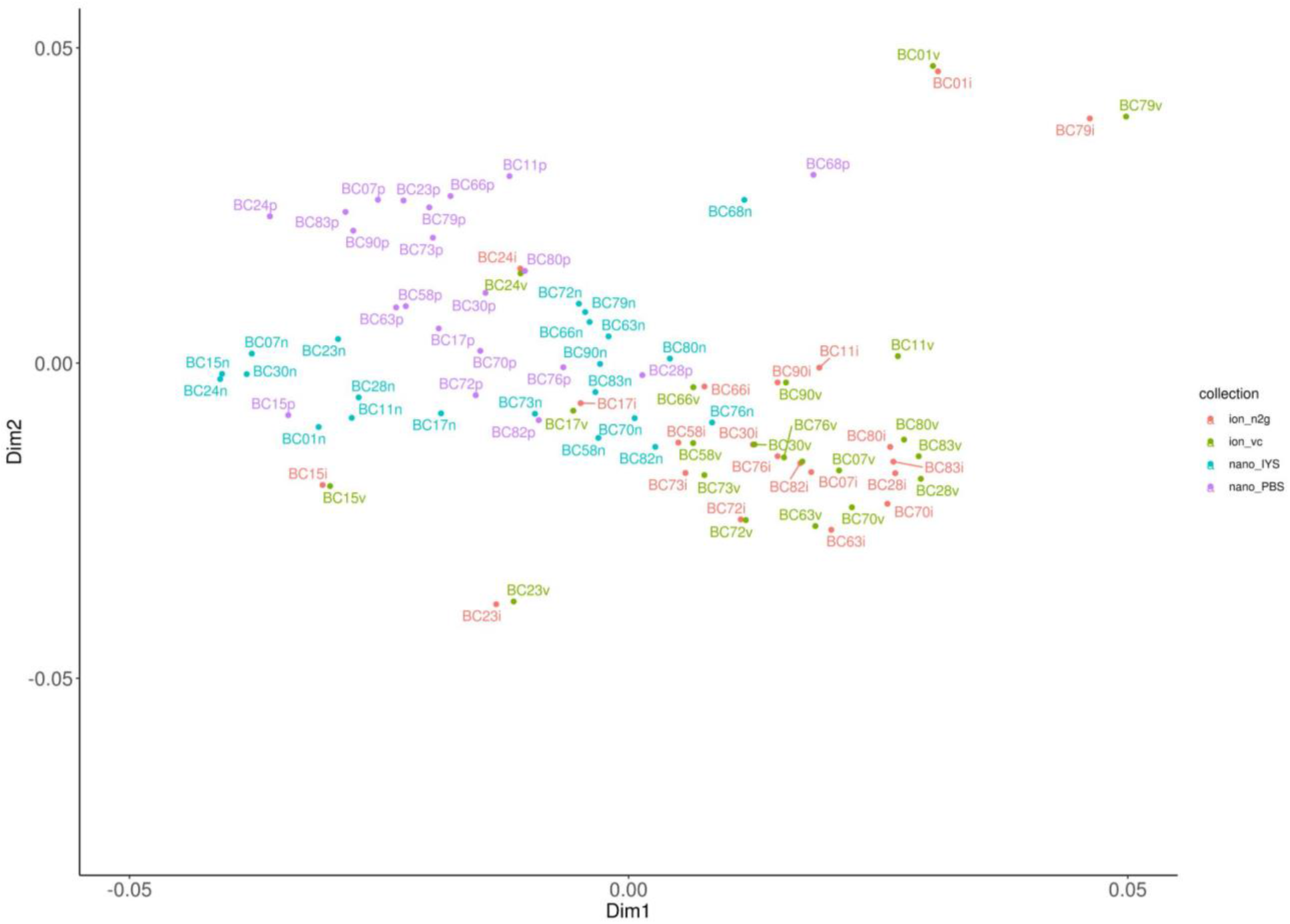
Principal coordinate analysis (PCoA) of SNP calls of individuals passing threshold in all datasets. SNP calls based on Nanopore sequences generated during the IYS expedition shown in blue (“nano_IYS”), and the same individuals reanalyzed upon return using the same workflow shown in purple (“nano_PBS”). Ion Torrent reads scored with the n2g pipeline are shown in red (“ion_n2g”) and scores derived from the Ion Torrent variant caller are shown in green (“ion_vc”).

**Figure 9:**
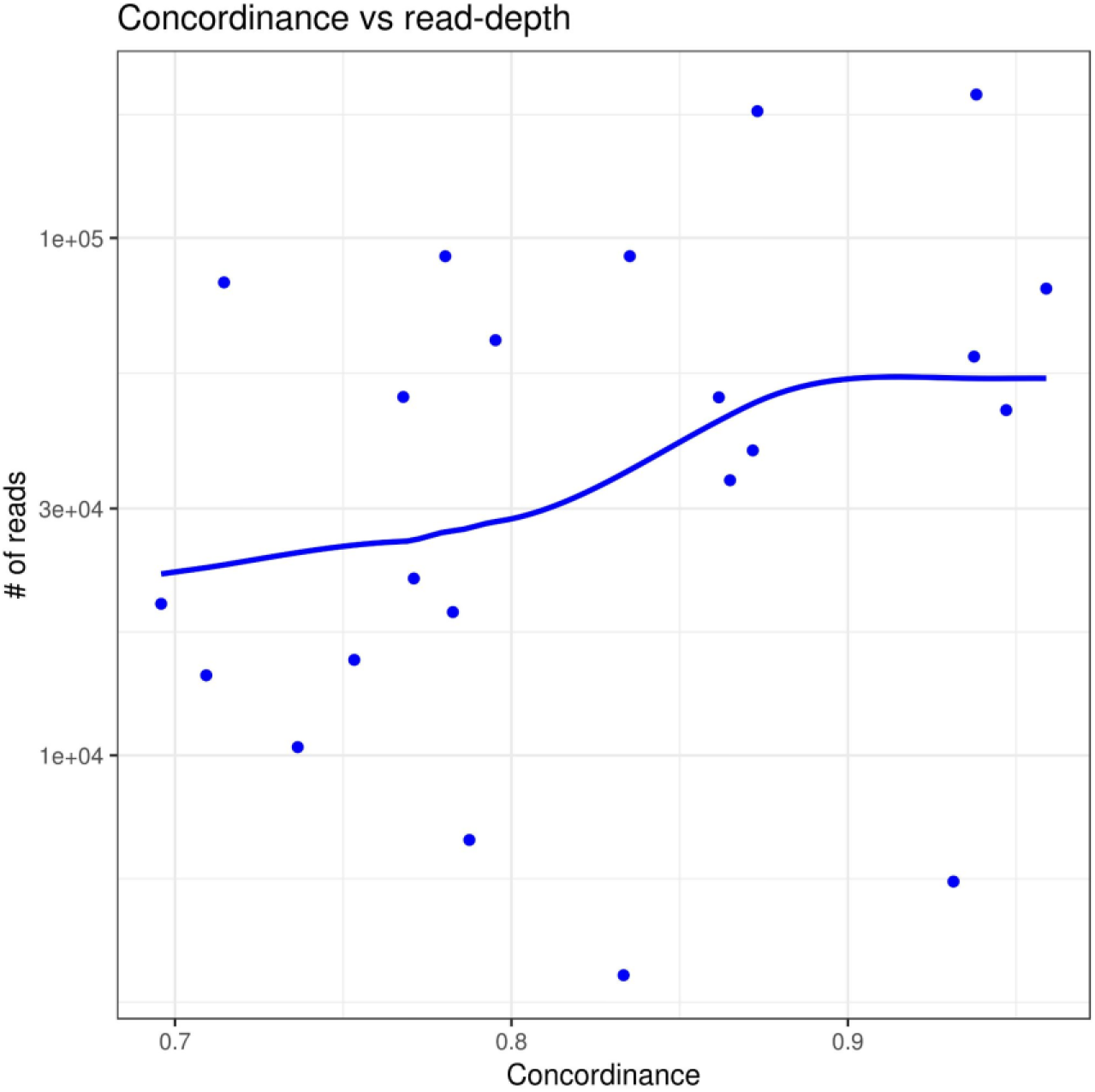
SNP call concordance between sequencing platforms shows weak correlation with Nanopore read counts. Concordance of Nanopore read based SNP calls with SNP calls generated from Ion Torrent based sequencing are plotted against read number in the associated barcode bin. Linear regression of the data by loess is depicted in lines corresponding in color with the samples.

### Stock assignment based on Nanopore data differs inherently from Ion Torrent based assignments in a subset of individuals

Stock assignment by rubias showed discrepancies between the Nanopore and Ion Torrent based datasets. In only 61.5% of cases did Nanopore sequences (PBS run) lead to the same top reporting unit (repunit; large scale geographic areas such as Westcoast Vancouver Island or Lower Fraser River) assignment for individual stock ID as the Ion Torrent based sequences (Figure 10, Table 1). Specifically, Nanopore-based repunit assignment showed higher proportions of assignments to South Eastern Alaska (SEAK) than Ion Torrent-based assignments (Figure 10, Table 1).

**Figure 10:**
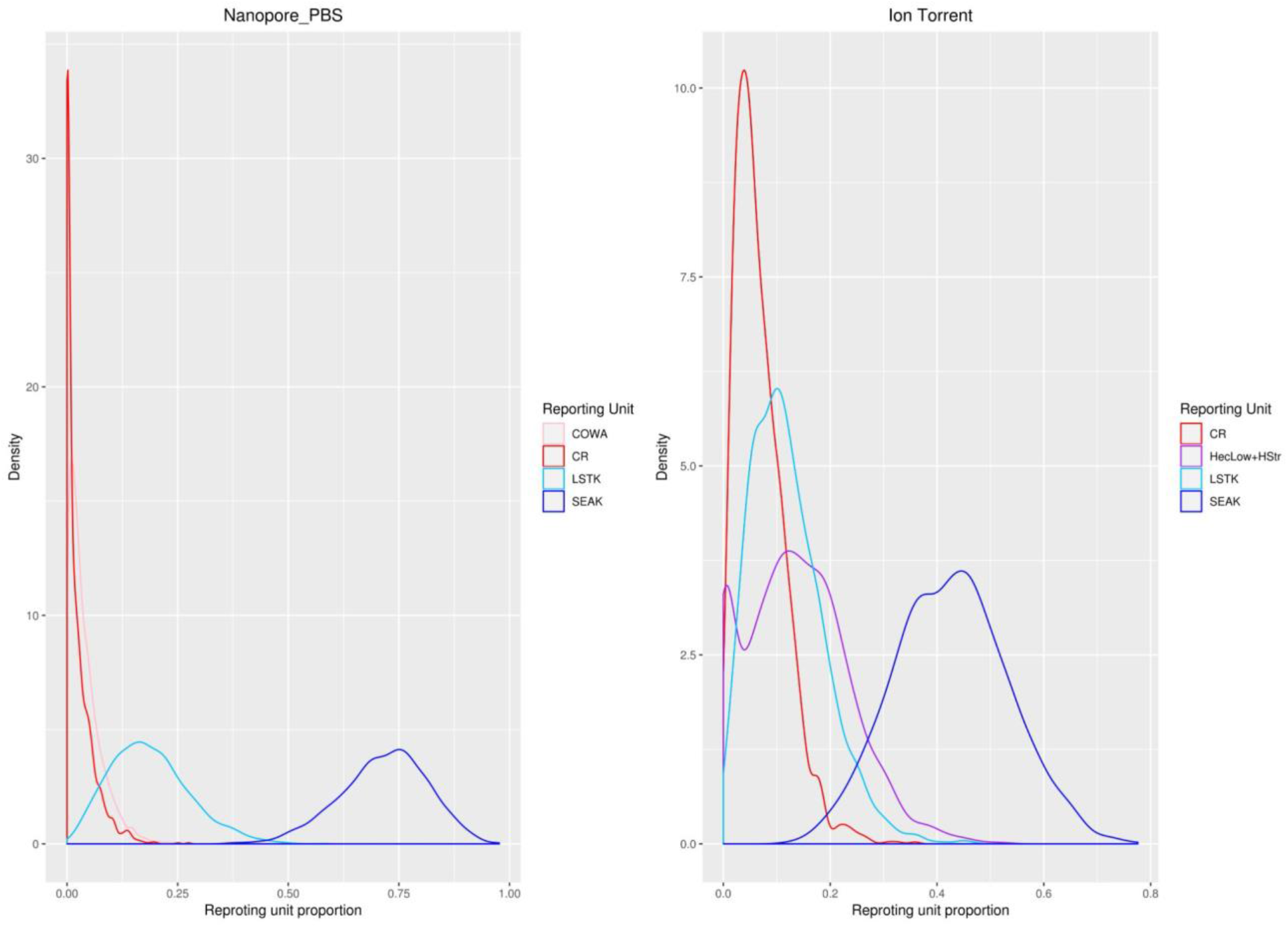
Relative proportion of reporting units to the overall mixture of coho salmon. Only individuals that had passed the stock ID threshold (>50% of SNPs called) on all three GSI runs are included. Reporting Units: SEAK: Southeast Alaska; LSTK: Lower Stikine River; HecLow+HStr: Lower Hecate Strait and Haro Strait; SC + SFj: Southern Coastal Streams, Queen Charlotte Strait, Johnston Strait and Southern Fjords; CR: Columbia River; COWA: Coastal Washington.

**Table1:** Relative proportion of top reporting units (contribution >3%) to the overall mixture of coho salmon. Only individuals that had successful stock ID on all three GSI runs are included. Reporting Units: SEAK: Southeast Alaska; LSTK: Lower Stikine River; NCS: North Coast Streams (BC); HecLow+HStr: Lower Hecate Strait and Haro Strait; SC + SFj: Southern Coastal Streams, Queen Charlotte Strait, Johnston Strait and Southern Fjords; CR: Columbia River; COWA: Coastal Washington; LNASS: Lower Nass River; WVI: West Vancouver Island; OR: Oregon.

|  | Ion Torrent (ion_vc) |  |  | Nanopore (nano_PBS) |  |  |
| --- | --- | --- | --- | --- | --- | --- |
| Rank | Repunit | Proportion | SD | Repunit | repprop | SD |
| 1 | SEAK | 0.437678 | 0.109758 | SEAK | 0.662083 | 0.218561 |
| 2 | HecLow+HStr | 0.178637 | 0.057264 | LSTK | 0.205116 | NA |
| 3 | LSTK | 0.068878 | NA | CR | 0.050276 | 0.012993 |
| 4 | SC+SFj | 0.067989 | 0.025318 | COWA | 0.042244 | 0.011583 |
| 5 | CR | 0.067939 | 0.01403 |  |  |  |
| 6 | NCS | 0.036009 | 0.004052 |  |  |  |
| 7 | OR | 0.034352 | 0.010704 |  |  |  |
| 8 | WVI | 0.033487 | 0.009144 |  |  |  |
| 9 | LNASS | 0.032288 | 0.022742 |  |  |  |

Nevertheless, mixture proportions in both datasets were dominated by South Eastern Alaska stocks. Nanopore assignments tendedto overestimate the contribution to this stock as well as Lower Stikine River stocks (LSTK). Many of the individuals assigned to these stocks using the Nanopore were assigned to the adjacent stocks of Lower Hecate Strait and Haro Strait (HecLow+HStr) as well as Southern Coastal Streams, Queen Charlotte Strait, Johnston Strait and Southern Fjords (SC + SFj) on the Ion Torrent platform (Figure 10, Table 1). Individuals from stocks well represented in the database like the Columbia River were confidently assigned to the appropriate stock on both platforms.

However, Z-scores calculated by rubias during stock assignment, which are an indirect measure of how well the SNP call match individuals in the baseline dataset of both, indicated that the Nanopore and the Ion Torrent data showed large deviations from the normal distribution, suggesting that many individuals assayed are not well represented in the database (Figure 11) (Moran and Anderson 2019). Ion Torrent data shows two peaks, one overlaying the expected normal distribution and a second peak that lay outside of the normal distribution. This suggests that about half of the individuals were not from populations that are well represented in the database (Figure 11). Similarly, Nanopore-based assignments showed even more aberrant distribution, presumably due to the additive effects of the sequencing platform introducing bias on top of poor baseline representation (Figure 11). The poor database representation could cause small differences in SNP calls to cause alternative assignments.

**Figure 11:**
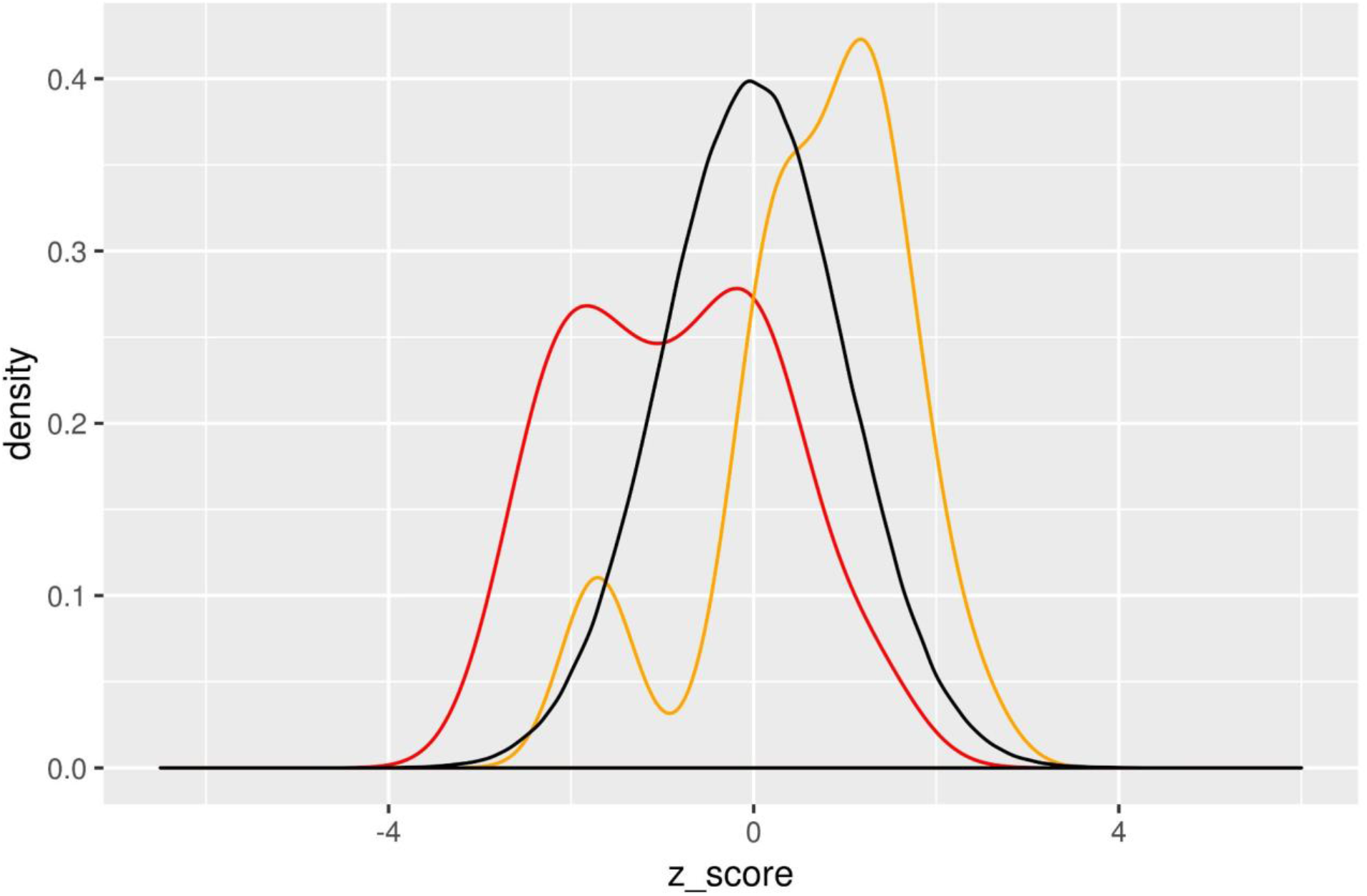
Z-score distribution of reference assignment by sequencing run: Black: Normal distribution, Red: Ion Torrent data, Yellow: PBS Nanopore data. A perfect representation of the assessed individuals would overlap the normal distribution.

## Discussion

### Nanopore sequencing enables remote in-field single nucleotide polymorphism genetic stock identification

Here, we present the first proof-of-concept study demonstrating the feasibility of using the portable Oxford Nanopore minION sequencer for remote in-field genetic stock identification by SNP sequencing of Pacific salmon. We developed a rapid sample processing workflow that relied on amplicon concatenation to increase throughput. With this workflow, we performed genetic stock identification on 75 coho salmon onboard a research vessel in the Gulf of Alaska, with minimal equipment during two runs. Genetic stock identification of all 80 captured coho salmon in a single run using the mobile platform resulted in stock assignment for 50 individuals at 67% concordance with state of the art laboratory based pipelines.

Despite its promising performance, the fidelity, throughput, and turnaround time of Nanopore-based SNP GSI currently still falls short of what would enable this technology to be used for the wide range of remote real-time applications we intended it for. This is due to a number of factors, such as inefficient barcoding, error rates, inefficiencies of custom genotyping pipelines, low concatenation efficiency, and limited computational power in our setup.

The inherent low fidelity of the Nanopore platform using R9 type flow cells relative to other sequencing technologies, specifically around homopolymer tracts, proved to be the major shortcoming, limiting both the actual SNP calling accuracy, causing comparatively low repeatability, as well as the throughput, by necessitating a higher alignment coverage due to the high error rate (Cornelis et al. 2017). The low fidelity of the Nanopore sequences was specifically apparent when comparing it with the established sequencing platform for genetic stock identification by SNP sequencing, the Ion Torrent Proton sequencer (Beacham et al. 2017). The Ion Torrent short read sequencer routinely outperformed the Nanopore sequencer, both in accuracy and in throughput. The latter being a major restricting factor of the Nanopore platform due to a limited number of available sequencing pores inherent to the platform. While we compensated for this limitation by concatenating amplicons, to generate several amplicon sequences per Nanopore read, the efficiency of this approach was modest, yielding only a two-fold increase in throughput at present. Further, the needs for concatenation and higher inputs required several PCR amplification steps that could have contributed to the observed shifts in allele frequencies leading to differing assignments on the different platforms. Turnaround time in the present study was mostly restricted by the computational capacity of the portable laptop used for the computational analysis. Specifically, base calling by translating the raw electrical signal recorded by the minION sequencer into fastq nucleotide reads proved to be the most time-consuming step, requiring up to several days in computing time.

However, despite the limitations associated with the Nanopore platform described above, the stock composition of coho in the Gulf of Alaska also confounded accuracy and fidelity of stock assignment. Most importantly, the majority of fish sampled and assessed during the Gulf of Alaska expedition were assigned to Southeastern Alaska and adjacent British Columbina coast stocks (SEAK, HecLow+HStr, SC + SFj). These stocks are poorly represented in the queried baseline and stocks from northern Alaska are very sparse so that fish from such origin often get assigned to the SEAK with poor confidence. This meant that even on the Ion Torrent platform, assignment probabilities were low, causing small differences in SNP content between the two platforms to lead to alternating assignment between these stocks (i.e. SEAK assignment on Nanaopore being assigned to HecLow+HStr and SC + SFj on Ion Torrent). Indeed, stock assignment on the Ion Torrent platform using an updated and expanded baseline and primer set, resulted in high confidence assignment of many of these individuals to Kynoch and Mussel Inlets, a spatially close reporting unit on the Northern BC coast that was poorly represented in the original baseline (C. Neville, personal communication). This suggests that new loci included in the updated primer set and baseline were able to resolve these stocks at higher confidence and assign them to the appropriate stock (Beacham et al. 2020). Fortunately, all of the current limitations mentioned above will be addressed in further development and we expect significant improvements in all fields, ultimately delivering a high throughput, real-time, in-field sequencing platform.

### Advances to the Nanopore platform, sample preparation, as well as computational infrastructure will improve turnaround, throughput, and fidelity

While we were successful in providing a proof-of-principle study demonstrating that the Nanopore platform is capable of in-field genotyping, the throughput, fidelity, and turnaround, remained below the level needed to put this platform into standard operation for GSI by SNP genotyping. Several modifications in the workflow are planned to improve the throughput. Currently, barcoding relies on inefficient blunt-end ligation of the barcoding adapters to the PCR amplicons, leading to up to 50% unbarcoded amplicons and therefore wasting a large portion of sequencing capacity. Including the ligation adapter sequences needed to add the barcodes in the PCR primers will improve the efficacy of barcoding by circumventing the inefficient and laborious blunt-end ligation. This will improve sequencing throughput, while at the same time speeding up the sample preparation by approximately one hour. Next, concatenation efficiency is currently relatively low, increasing throughput only two-fold. While large concatemers approaching 10kb were observed, they were relatively rare. Optimized concatenation conditions by adjusting the reaction conditions such as annealing temperature and duration should exponentially improve throughput by both increasing the relative abundance of concatenated amplicons, as well as the total length of concatemers. Further workflow improvements could include pre-aliquoting of DNA extraction solution, barcodes and primers, as well as bead cleaning materials in 96 well plates before heading into the field, which should reduce an additional two hours of sample preparation, as well as reduce the risk of cross-contamination in the field. Together, these improvements should bring the total sample preparation time to about 10h, with approximately half the time being hands-on.

The major current bottleneck in turnaround time is the time that base calling takes on the portable laptop computer used in the present study. The Nanopore computation unit minIT, however, can provide real-time base calling to fastq and is currently being tested in the follow-up work to the present study. Actual real-time basecalling will bring the workflow in the neighbourhood of the desired 24h turnaround time.

An additional issue for using Nanopore sequencing is the low accuracy of the sequencing platform at the time of this project using the R9 flow cells. This low accuracy requires excessively high alignment coverage at SNP locations to ensure accurate SNP calling. However, newer Nanopore flow cells promise greatly increased accuracy (e.g., 99.999% for R10) due to “a longer barrel and dual reader head” and have recently become available. This updated flow cell technology is therefore expected to greatly improve sequencing accuracy and possibly allow the lowering of alignment thresholds for SNP calling, thereby increasing the throughput more than twofold. Improvements to the SNP calling pipeline, might enable the identification and exclusion of erroneous SNP calls due to the ability to calculate the p-error associated with SNP calls, thereby increasing accuracy and repeatability. Finally, in selecting SNP loci for inclusion in GSI baselines, consideration of the types of sequences that are most problematic for Nanopore sequencing (e.g. homopolymer tracts) could go a long way to improving performance across platforms. Testing power in coastwide baselines once these problematic loci are excluded will be an important future step. Extrapolating the above mentioned improvements would improve the current throughput of 96 individuals per flow cell by more than an order of magnitude, thereby enabling cost-effective real-time and/or field-based application of the platform.

Currently, Nanopore-based SNP GSI is an experimental in-field stock identification tool. Turnaround of several days and throughput limited to only 96 individuals per flow cell limit its attractiveness for a wider user base. Future improvements of the sequencing platform, the sample preparation procedure as well as the computational infrastructure will greatly improve throughput and turnaround for this. This should enable the application of Nanopore-based SNP GSI for near-real-time stock management of variable batch sizes at-sea or in remote locations. Further, parallel sequencing on several flow cells using the Oxford Nanopore GridION, which can employ five flow cells simultaneously, would enable dynamic real-time stock identification using variable batch sizes from dozens to hundreds of individuals. In the event that rapid turnaround is required, the sequencing library can also be spread across several flow cells on the GridION. Together, these updates would greatly improve the abilities of multiple user groups including government, Indigenous communities, and conservation organizations to conduct GSI for safeguarding populations at risk, while allowing sustainable harvest of healthy populations.

## Acknowledgements

The authors would like to thank the following individuals for their contribution to the expedition and to the manuscript: Richard Beamish, Brian Riddell, and the NPAFC secretariat for the organization of the 2019 Gulf of Alaska expedition. The entire scientific crew of the 2019 GoA expedition: Evgeny Pakhomov, Gerard Foley, Brian P.V. Hunt, Arkadii Ivanov, Hae Kun Jung, Gennady Kantakov, Anton Khleborodov, Chrys Neville, Vladimir Radchenko, Igor Shurpa, Alexander Slabinsky, Shigehiko Urawa, Anna Vazhova, Vishnu Suseelan, Charles Waters, Laurie Weitkamp, and Mikhail Zuev. The crew of the research vessel Professor Kaganovskiy. Charles Waters for providing an R script for catch visualization. Chrys Neville for the contribution of catch data. This research was supported by Pacific Salmon Commission, Pacific Salmon Foundation, and Fisheries and Oceans Canada and the Canadian Coast Guard (DFO CCG). CMD was supported by a fellowship through the Pacific Salmon Foundation and MITACS.

## Supplementary materials

**Supplementary Table 1:**
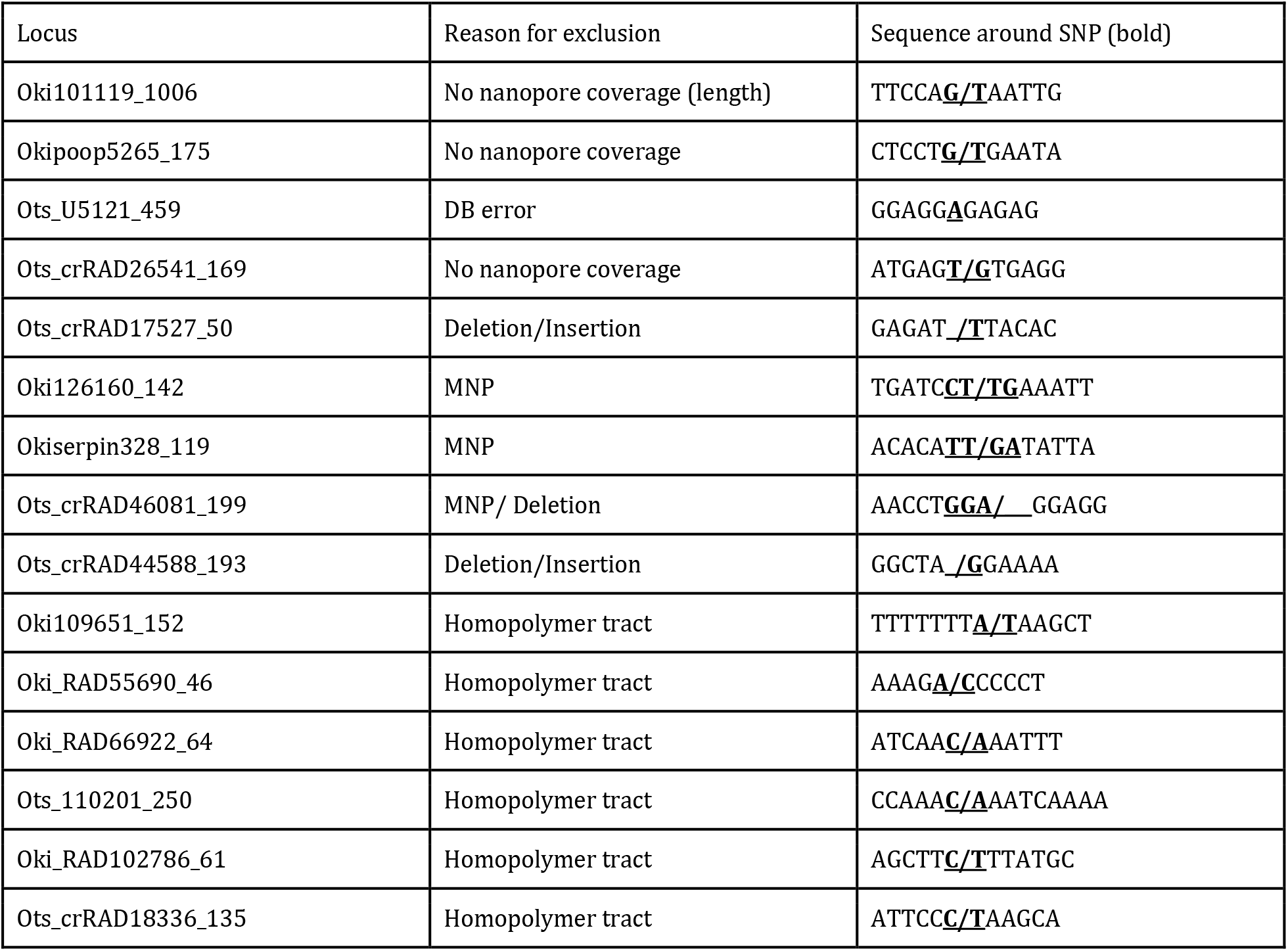
SNP loci excluded from the nanopore analysis

**Supplementary Table 2:** Representation of individuals in the queried baseline.

| *Reporting Unit* | *Abbreviation* | *Individuals* |
| --- | --- | --- |
| *Russia* | *Russia* | *357* |
| *Southeast Alaska* | *SEAK* | *814* |
| *Alsek River* | *ALSEK* | *96* |
| *Lower Stikine* | *LSTK* | *40* |
| *Portland Sound-Observatory Inlet-Portland Canal* | *PORT* | *99* |
| *Lower Nass* | *LNASS* | *191* |
| *Upper Nass* | *UNASS* | *92* |
| *Lower Skeena* | *LSKNA* | *287* |
| *Skeena Estuary* | *SKEst* | *119* |
| *Middle Skeena* | *MSKNA* | *269* |
| *Upper Skeena* | *USKNA* | *244* |
| *Haida Gwaii-Graham Island Lowlands* | *NHG* | *467* |
| *Haida Gwaii-East* | *EHG* | *315* |
| *Haida Gwaii-West* | *WHG* | *136* |
| *Northern Coastal Streams* | *NCS* | *1307* |
| *Hecate Strait Mainland* | *HecLow+HStr* | *409* |
| *Mussel-Kynoch* | *MusKyn* | *182* |
| *Douglas Channel-Kitimat Arm* | *DOUG* | *476* |
| *Bella Coola-Dean Rivers* | *BCD* | *348* |
| *Rivers Inlet* | *Rivers* | *368* |
| *Smith Inlet* | *Smith* | *212* |
| *Southern Coastal Streams-Queen Charlotte Strait-Johnstone Strait-Southern Fjords* | *SC+SFj* | *599* |
| *Homathko-Klinaklini Rivers* | *HK* | *174* |
| *Georgia Strait Mainland* | *SC+GStr* | *162* |
| *Howe Sound-Burrard Inlet* | *Howe-Burrard* | *5093* |
| *Nahwitti Lowland* | *Nahwitti* | *993* |
| *East Vancouver Island-Georgia Strait* | *EVI+GStr* | *9221* |
| *East Vancouver Island-Johnstone Strait-Southern Fjords* | *EVI+SFj* | *49* |
| *West Vancouver Island* | *WVI* | *2079* |
| *Clayoquot* | *CLAY* | *343* |
| *Juan de Fuca-Pachena* | *JdF* | *2310* |
| *Upper Fraser* | *UFR* | *45* |
| *Lower Fraser* | *LFR* | *10368* |
| *Lillooet* | *LILL* | *235* |
| *Fraser Canyon* | *FRCany* | *175* |
| *Interior Fraser* | *IntrFR* | *288* |
| *Lower Thompson* | *LTHOM* | *589* |
| *North Thompson* | *NTHOM* | *1276* |
| *South Thompson* | *STHOM* | *1509* |
| *Boundary Bay* | *BB* | *591* |
| *Northern Puget Sound* | *NPS* | *296* |
| *Mid-Puget Sound* | *MPS* | *233* |
| *Skagit River* | *SKAG* | *277* |
| *Southern Puget Sound* | *SPS* | *272* |
| *Juan de Fuca Strait* | *JUAN* | *145* |
| *Northern Washington* | *NOWA* | *196* |
| *Hood Canal* | *HOOD* | *167* |
| *Coastal Washington* | *COWA* | *369* |
| *Columbia River* | *CR* | *608* |
| *Oregon* | *OR* | *511* |

## References

Beacham, Terry D., Colin G. Wallace, Kim Jonsen, Brenda McIntosh, John R. Candy, Eric B. Rondeau, Jean-Sébastien Moore, Louis Bernatchez, and Ruth E. Withler. 2020. “Accurate Estimation of Conservation Unit Contribution to Coho Salmon Mixed-Stock Fisheries in British Columbia, Canada Using Direct DNA Sequencing for Single Nucleotide Polymorphisms.” Canadian Journal of Fisheries and Aquatic Sciences. Journal Canadien Des Sciences Halieutiques et Aquatiques, no. ja. https://www.nrcresearchpress.com/doi/abs/10.1139/cjfas-2019-0339.

Beacham, Terry D., Colin Wallace, Cathy MacConnachie, Kim Jonsen, Brenda McIntosh, John R. Candy, Robert H. Devlin, and Ruth E. Withler. 2017. “Population and Individual Identification of Coho Salmon in British Columbia through Parentage-Based Tagging and Genetic Stock Identification: An Alternative to Coded-Wire Tags.” Canadian Journal of Fisheries and Aquatic Sciences. Journal Canadien Des Sciences Halieutiques et Aquatiques 74 (9): 1391–1410.

Beacham, Terry D., Colin Wallace, Cathy MacConnachie, Kim Jonsen, Brenda McIntosh, John R. Candy, and Ruth E. Withler. 2018. “Population and Individual Identification of Chinook Salmon in British Columbia through Parentage-Based Tagging and Genetic Stock Identification with Single Nucleotide Polymorphisms.” Canadian Journal of Fisheries and Aquatic Sciences. Journal Canadien Des Sciences Halieutiques et Aquatiques 75 (7): 1096–1105.

Campbell, Nathan R., Stephanie A. Harmon, and Shawn R. Narum. 2015. “Genotyping-in-Thousands by Sequencing (GT-Seq): A Cost Effective SNP Genotyping Method Based on Custom Amplicon Sequencing.” Molecular Ecology Resources 15 (4): 855–67.

Cederholm, C. Jeff, Matt D. Kunze, Takeshi Murota, and Atuhiro Sibatani. 1999. “Pacific Salmon Carcasses: Essential Contributions of Nutrients and Energy for Aquatic and Terrestrial Ecosystems.” Fisheries 24 (10): 6–15.

Cook, Rodney C., and I. Guthrie. 1987. “In-Season Stock Identification of Sockeye Salmon (Oncorhynchus Nerka) Using Scale Pattern Recognition.” Canadian Special Publication of Fisheries and Aquatic sciences/Publication Speciale Canadienne Des Sciences Halieutiques et Aquatiques 96: 327–34.

Cornelis, Senne, Yannick Gansemans, Lieselot Deleye, Dieter Deforce, and Filip Van Nieuwerburgh. 2017. “Forensic SNP Genotyping Using Nanopore MinION Sequencing.” Scientific Reports 7 (February): 41759.

Gilbey, John, Vidar Wennevik, Ian R. Bradbury, Peder Fiske, Lars Petter Hansen, Jan Arge Jacobsen, and Ted Potter. 2017. “Genetic Stock Identification of Atlantic Salmon Caught in the Faroese Fishery.” Fisheries Research 187 (March): 110–19.

Hinch, S. G., S. J. Cooke, A. P. Farrell, K. M. Miller, M. Lapointe, and D. A. Patterson. 2012. “Dead Fish Swimming: A Review of Research on the Early Migration and High Premature Mortality in Adult Fraser River Sockeye Salmon Oncorhynchus Nerka.” Journal of Fish Biology 81 (2): 576–99.

Holtby, L. Blair, Bruce C. Andersen, and Ronald K. Kadowaki. 1990. “Importance of Smolt Size and Early Ocean Growth to Interannual Variability in Marine Survival of Coho Salmon (Oncorhynchus Kisutch).” Canadian Journal of Fisheries and Aquatic Sciences. Journal Canadien Des Sciences Halieutiques et Aquatiques 47 (11): 2181–94.

Jefferts, K. B., P. K. Bergman, and H. F. Fiscus. 1963. “A Coded Wire Identification System for Macro–Organisms.” Nature 198 (4879): 460–62.

Lichatowich, Jim. 2001. Salmon Without Rivers: A History Of The Pacific Salmon Crisis. Island Press.

Li, Heng, and Richard Durbin. 2009. “Fast and Accurate Short Read Alignment with Burrows-Wheeler Transform.” Bioinformatics 25 (14): 1754–60.

Li, Heng, Bob Handsaker, Alec Wysoker, Tim Fennell, Jue Ruan, Nils Homer, Gabor Marth, Goncalo Abecasis, Richard Durbin, and 1000 Genome Project Data Processing Subgroup. 2009. “The Sequence Alignment/Map Format and SAMtools.” Bioinformatics 25 (16): 2078–79.

Mikheyev, Alexander S., and Mandy M. Y. Tin. 2014. “A First Look at the Oxford Nanopore MinION Sequencer.” Molecular Ecology Resources 14 (6): 1097–1102.

Miller, Kristina M., Amy Teffer, Strahan Tucker, Shaorong Li, Angela D. Schulze, Marc Trudel, Francis Juanes, et al. 2014. “Infectious Disease, Shifting Climates, and Opportunistic Predators: Cumulative Factors Potentially Impacting Wild Salmon Declines.” Evolutionary Applications 7 (7): 812–55.

Miller, Kristina M., Ruth E. Withler, and Terry D. Beacham. 1996. “Stock Identification of Coho Salmon (Oncorhynchus Kisutch) Using Minisatellite DNA Variation.” Canadian Journal of Fisheries and Aquatic Sciences. Journal Canadien Des Sciences Halieutiques et Aquatiques 53 (1): 181–95.

Moran, Benjamin M., and Eric C. Anderson. 2019. “Bayesian Inference from the Conditional Genetic Stock Identification Model.” Canadian Journal of Fisheries and Aquatic Sciences. Journal Canadien Des Sciences Halieutiques et Aquatiques 76 (4): 551–60.

Ozerov, Mikhail, Anti Vasemägi, Vidar Wennevik, Rogelio Diaz-Fernandez, Matthew Kent, John Gilbey, Sergey Prusov, Eero Niemelä, and Juha-Pekka Vähä. 2013. “Finding Markers That Make a Difference: DNA Pooling and SNP-Arrays Identify Population Informative Markers for Genetic Stock Identification.” PloS One 8 (12): e82434.

Paradis, Emmanuel, and Klaus Schliep. 2019. “Ape 5.0: An Environment for Modern Phylogenetics and Evolutionary Analyses in R.” Bioinformatics 35 (3): 526–28.

Quick, Joshua, Nicholas J. Loman, Sophie Duraffour, Jared T. Simpson, Ettore Severi, Lauren Cowley, Joseph Akoi Bore, et al. 2016. “Real-Time, Portable Genome Sequencing for Ebola Surveillance.” Nature 530 (7589): 228–32.

Robinson, James T., Helga Thorvaldsdóttir, Wendy Winckler, Mitchell Guttman, Eric S. Lander, Gad Getz, and Jill P. Mesirov. 2011. “Integrative Genomics Viewer.” Nature Biotechnology 29 (1): 24–26.

Schlecht, Ulrich, Janine Mok, Carolina Dallett, and Jan Berka. 2017. “ConcatSeq: A Method for Increasing Throughput of Single Molecule Sequencing by Concatenating Short DNA Fragments.” Scientific Reports 7 (1): 5252.

Wood, Chris C., Dennis T. Rutherford, and Skip McKinnell. 1989. “Identification of Sockeye Salmon (Oncorhynehus Nerka) Stocks in Mixed-Stock Fisheries in British Columbia and Southeast Alaska Using Biological Markers.” Canadian Journal of Fisheries and Aquatic Sciences. Journal Canadien Des Sciences Halieutiques et Aquatiques 46 (12): 2108–20.

Woodey, J. C. 1987. “In-Season Management of Fraser River Sockeye Salmon (Oncorhynchus Nerka): Meeting Multiple Objectives.” Sockeye Salmon, 367–74.

